# Integrated biophysical and spatial remodeling during insulin secretory granule maturation at the mitochondrial network

**DOI:** 10.64898/2026.03.02.709121

**Authors:** Rachel Knight, Aneesh Deshmukh, Wen Lin, Riva Verma, Kate White

## Abstract

Effective insulin secretion and blood glucose homeostasis depend on the multistep maturation of insulin secretory granules (ISGs), a process that includes lumen acidification, enzymatic insulin processing, and biophysical remodeling of the granule. An under studied aspect of ISG maturation is the role of inter-organelle contacts in organelle remodeling. While a correlation between ISG-mitochondria contacts and ISG maturation has been observed, many questions remain on how this interaction may impact maturation (1–5). We sought to address this gap in knowledge by using multi-scale imaging approaches (fluorescent microscopy, soft X-ray tomography, and cryo-electron tomography) to examine how the biophysical properties and spatial organization of ISGs change around the mitochondrial network. Our data suggests that ISGs in proximity to mitochondria exhibit lower pH, higher biomolecular density, and smaller vesicle diameter. Time-resolved imaging using a SNAP tag labelling system also shows that as ISGs age, their proximity to the mitochondria network is increased between 3-6 hours after biosynthesis, suggesting that ISG-mitochondria association is dynamically spatiotemporally regulated in pancreatic β-cells. These data suggest that mitochondrial proximity contributes to the maturation and remodeling of ISGs in pancreatic beta cells.

## Introduction

Insulin is a vital peptide hormone secreted from pancreatic β-cells to maintain blood glucose homeostasis (6, 7). Insulin is processed, stored, transported, and secreted in insulin secretory granules (ISGs) (6, 8). ISGs undergo a maturation process that includes the acidification of the granule lumen and enzymatic processing of proinsulin to active insulin, condensation of insulin into a dense core or crystal within the lumen, and lipid and protein remodeling of the ISG. (6, 8–10). Defects in ISG maturation and ISG lipid and protein content have been implicated in Type 1 and Type 2 diabetes, suggesting a possible role in disease by which ISG maturation contributes to disease (11–14).

A key gap in knowledge is how ISG maturation and remodeling are regulated within the cell. In addition to enzymatic processing, maturation involves dynamic remodeling of granule membrane composition and biophysical properties. There is precedent for ISG lipid exchange occurring via inter-organelle contacts, specifically interactions between ISGs and the Endoplasmic Reticulum (15). Suggesting that inter-organelle contacts can play a role in ISG biophysical remodeling.

Membrane contact sites have broadly been observed for decades with marked functions in signaling, lipid transfer, and membrane dynamics (16, 17). Many well-characterized contact sites revolve around the mitochondrial network, facilitating lipid and metabolite transfer with other organelles, such as the ER (15, 18–20). Although ISGs are frequently observed in proximity to or contacting mitochondria (21, 22), the functional significance of these interactions remains largely unexplored. Specifically, it is unclear when during the ISG lifecycle these contacts occur, how long they persist, and which biophysical properties of the granule are influenced by mitochondrial association.

To characterize the relationship of mitochondria-ISG contact sites, we leveraged live-cell and high-resolution imaging approaches that provide complementary spatial and structural information across scales. These studies were performed primarily in INS-1E cells, an immortalized rat pancreatic β-cell line widely used to model insulin secretion, with select ultrastructural analyses conducted in primary mouse β-cells. Cryo-electron tomography (Cryo-ET) and soft X-ray tomography (SXT) provide high-resolution detail of the contact site in nearly native cells at sub-nanometer and nanometer resolution, respectively(21, 22). Further, the use of SXT also allows for direct quantification of ISG maturation state through changes in biomolecular density as mature ISG contain dense core or crystalline insulin (22–25). Fluorescence-based methods allow for mapping more dynamic changes of ISG trafficking and biochemical state. Fluorescence lifetime imaging microscopy (FLIM) based pH imaging provides insight into chemical state of ISGs in live cells within the context of the mitochondrial network. Time-resolved SNAP-tag based pulse-chase imaging methods allow for spatial mapping of ISGs at specific timepoints of their lifecycle relative to mitochondria.

Age has been shown to have multiple roles in the function of ISGs, with evidence indicating that younger granules are preferentially secreted in response to glucose stimulation, exhibit greater mobility, and maintain a lower oxidative state (26–30). These functional properties change as granules mature, suggesting that age can be used as a natural and unbiased metric to understand ISG dynamics. Accordingly, ISG age serves as an integrative measure of granule state and provides a contextual framework for interpreting maturation-associated readouts such as granule pH and molecular density. Using pulse-chase confocal imaging to temporally define granule age, we observed that mitochondrial proximity is not uniform across the ISG lifetime but is instead enriched during intermediate stages. Complementary FLIM-FRET measurements revealed that ISGs positioned near the mitochondrial network exhibit increased acidity, while SXT analysis demonstrated greater molecular density in these mitochondria-proximal granules, consistent with progressive insulin condensation. Together, these findings indicate that key maturation-associated changes, such as acidification and increased biomolecular density, may coincide with periods of enhanced mitochondrial proximity. Further, cryo-ET data in primary mouse β-cells suggest that these spatial relationships may be mediated through direct membrane contact sites, supporting a model in which physical interactions between ISGs and mitochondria facilitate metabolic exchange during defined windows of granule maturation.

## Results

### ISG-mitochondria associations can be observed across imaging scales

We first set out to image ISG-mitochondria contacts in live cells. Super-resolution time-lapse fluorescent imaging of a confined cellular region revealed dynamic interactions between ISGs and the mitochondrial network, in which individual ISGs were observed moving along the mitochondrial network (**Supplemental Video 1**). Notably, some ISGs transiently paused in close proximity to the mitochondria, indicating the formation of short-lived contacts lasting on the order of seconds. To place these dynamic observations in a broader cellular context, fixed cell fluorescent microscopy was used to label both ISG and mitochondria and observe ISG-mitochondria proximity at the whole cell level, revealing that labeled ISGs appear as discrete puncta dispersed along the mitochondrial network (**Fig. 1**). SXT was then used to further validate this observation in three-dimensional, fully hydrated cells. These tomograms demonstrate that individual granules are positioned immediately adjacent to mitochondria in 3D space. High-resolution cryo-ET imaging provided nanometer-scale snapshots of this interaction, validating that ISGs are positioned less than 10nm away from the mitochondrial membrane. These complementary imaging methods across scales establish that ISG-mitochondria contacts are ubiquitous and can be seen multiple times in each modality (**Figure S1**). The short distance observed between ISGs and mitochondria in our cryo-ET datasets places these organelles well within close enough proximity to facilitate the transfer of lipids or other metabolites essential for maturation. Such interactions may resemble established membrane contact sites, such as mitochondria–ER or ISG–ER interfaces, and could potentially facilitate ISG maturation. Together, these data establish a structural baseline, from whole cell scale organization to nanoscale proximity, providing a framework for testing whether ISG-mitochondria proximity advances insulin maturation.

**Figure 1.**
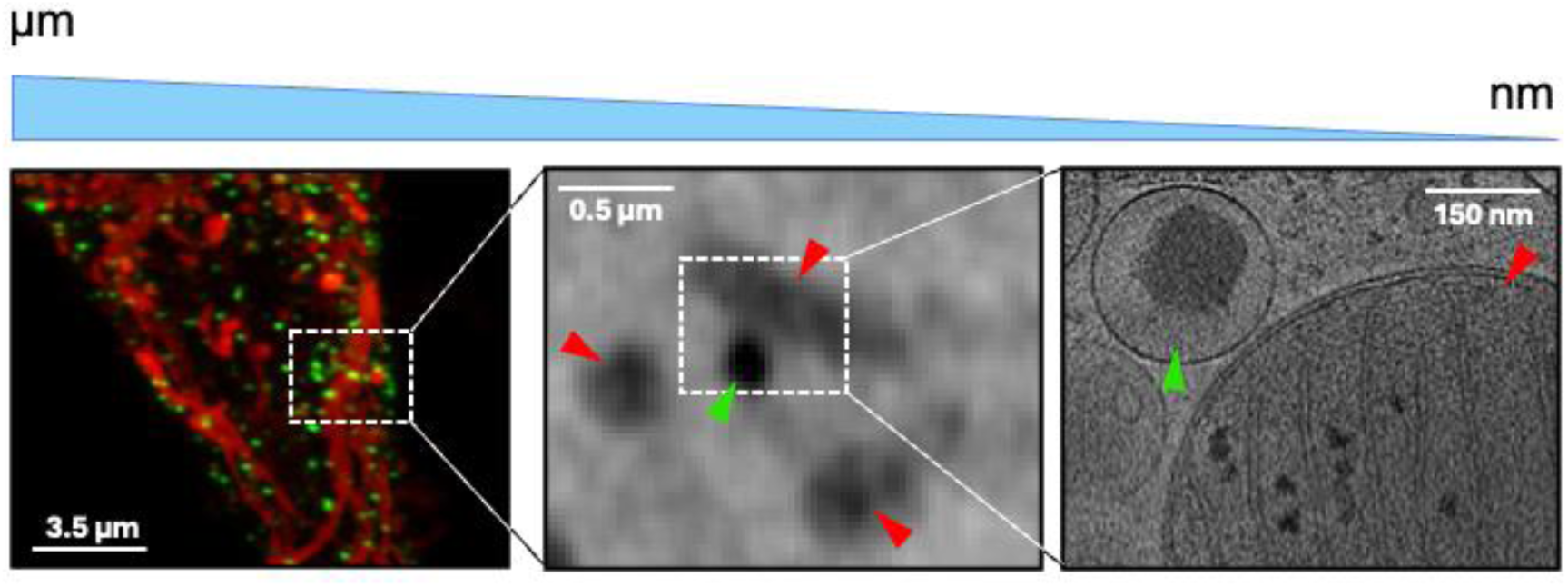
Correlative fluorescence and cryo-electron tomography reveal ISG-positive vesicles in close apposition to mitochondria. Left: Representative fluorescence micrograph showing mitochondria (red) and ISG-labeled puncta (green). The dashed box indicates the region selected for ultrastructural analysis. Scale bar, 3.5 µm. Middle: Soft X-ray tomographic slice of the highlighting electron-dense vesicular structures (red arrowheads) in proximity to a mitochondrion (green arrowhead). Scale bar, 0.5 µm. Right: Cryo-electron tomographic slice showing an ISG (green arrowhead) close to the outer mitochondrial membrane (red arrowhead). Scale bar, 150 nm.

### Soft X-Ray Tomography reveals biophysical characteristics of ISGs near mitochondria

Previous work has established the use of SXT to map ISG structure and biophysical characteristics in whole, fully hydrated cells at a 30 nm resolution (21, 23, 25). 3D segmentations of unstimulated INS-1E cells (starved for 30 minutes) revealed numerous ISGs distributed within the cell, alongside a dense mitochondrial network (**Figure 2A**). We used a 50 nm distance cut-off to define ISGs in “contact” with mitochondria, while all the other ISGs were classified as having “no contact” (**Figure 2B**). This threshold was selected based on the higher spatial resolution of SXT relative to fluorescence microscopy and falls within the range typically used to define membrane contact sites at the ultrastructural level. Importantly, although the absolute cutoff differs from that used in our fluorescence-based analyses due to modality-specific resolution constraints, the proportion of ISGs classified as mitochondria-associated remains consistent across imaging platforms. After pooling ISGs across cells, we next compared the properties of these two classes of ISGs. We observed that ISGs in contact with mitochondria were preferentially localized towards the cell interior (**Figure 2C**). When split into distinct Euclidean distance transform (EDT) zones, we observed a significant decrease in the raw distance between ISGs and mitochondria from normalized EDT zones 0-0.1 to 0.1-0.2 and 0.2-0.3. Beyond these zones, further reductions in ISG–mitochondria distance were not significant (**Figure 2D**).

**Figure 2:**
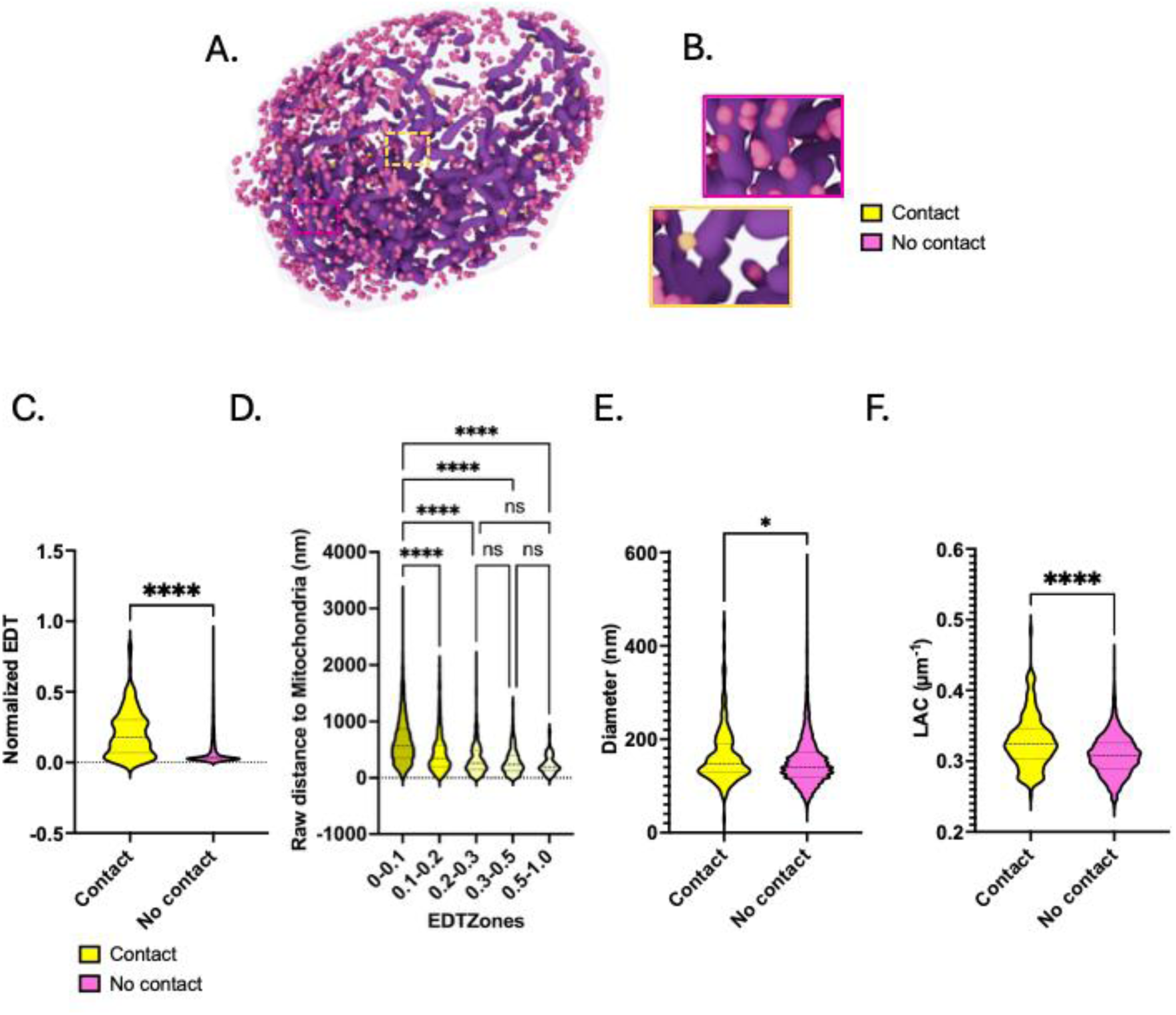
Soft X-Ray Tomography reveals biophysical characteristics of ISGs near mitochondria. (A) Representative whole-cell 3D model generated from a SXT tomogram of an unstimulated INS-1E cell showing the mitochondria network (purple), ISGs near the mitochondria (yellow), and distanced from the mitochondria (pink). Threshold for contact and no contact is 50nm. (B) Close-up views of the yellow and pink insets in (A) corresponding to ISGs in contact and having no-contact with the mitochondria respectively. (C) Comparison of the distance of the two types of ISGs from the plasma membrane. Normalized Euclidean distance transform values are used as a proxy for distance from the cell periphery. Comparison between ISGs in contact and not in contact with the mitochondria (D) Spatial distribution of ISGs based on distance from mitochondria. Distance to mitochondria progressively decreases towards the interior of the cell (higher EDT zones), with no significant difference between EDT range 0.2-1.0. *p<0.05; **p<0.01: ***p<0.001; ****p<0.0001 as calculated using a Welch’s t-test. (n = 186 for ISGs in contact with mitochondria, yellow; n = 5762 for ISGs not in contact with mitochondria (E) ISG diameters for ISGs either in contact or not in contact with the mitochondrial network show a significantly higher diameter for ISGs in contact with mitochondria. (F) Comparison between ISG biomolecular density (LAC value) for ISGs either in contact or not in contact with the mitochondrial network show a significantly higher LAC value for ISGs in contact with mitochondria

These two classes of ISGs also differ in their biomolecular density and morphological characteristics. ISGs in contact with mitochondria exhibited significantly higher mean linear absorption coefficient (LAC) values and larger diameters as compared to ISGs having no contact (**Figures 2E-F**). This increase in LAC near the mitochondrial network could suggest that mitochondria contribute to the maturation of insulin through transfer of ions, metabolites, or biomolecules that promote maturation within the ISG.

In addition to ISGs, mitochondrial biomolecular density showed correlation with ISG contacts, as demonstrated by the analysis of mitochondrial LAC from cell SXT tomograms (31, 32). To visualize this, we pseudo-colored each voxel of the mitochondrial network according to its LAC value, revealing rich local variation in biomolecular density within mitochondria, with distinct LAC hotspots visible (**Figure S2A, B**). Interestingly, when ISGs were overlaid onto the pseudo-colored mitochondrial LAC map, ISGs in contact with mitochondria localize to these LAC hotspots (**Fig S2C, D**), which could potentially suggest localized enrichment of biomolecules such as proteins or lipids at ISG-mitochondria contact sites.

### ISG acidity is impacted by mitochondrial association

Acidification of ISGs is necessary for effective enzymatic processing of proinsulin to insulin, thus making ISG pH a valuable indicator to assess ISG maturation state (33, 34). To evaluate the relationship between ISG acidity and their spatial proximity to the mitochondrial network, we expressed a fusion construct of a genetically encoded FLIM pH sensor E^2^GFP and known ISG marker neuropeptide Y (NPY) to report ISG pH in INS-1Ecells and co stained mitochondria. (35–37). (**Figure 3A**). Across 21 cells analyzed under basal conditions, ISGs proximal to mitochondria (-0.355µm – 0.05µm from mitochondria) exhibited a significantly lower pH than ISGs located farther away (0.05µm – 1.0µm away from mitochondria). (**Figure 3B**) This trend is also evident in the ensemble analysis of ISG pH on the phasor histogram: as granules shift toward the more acidic end of the distribution, their mean distance to mitochondria decreases significantly. Together, these results support the conclusion that mitochondria-proximal ISGs are more acidic. (**Figure 3C)** This data also shows a subpopulation of granules that are very acidic and not in contact with the mitochondrial network. We believe that these represent ISGs that are in recycling compartments or lysosomes for degradation.

**Figure 3.**
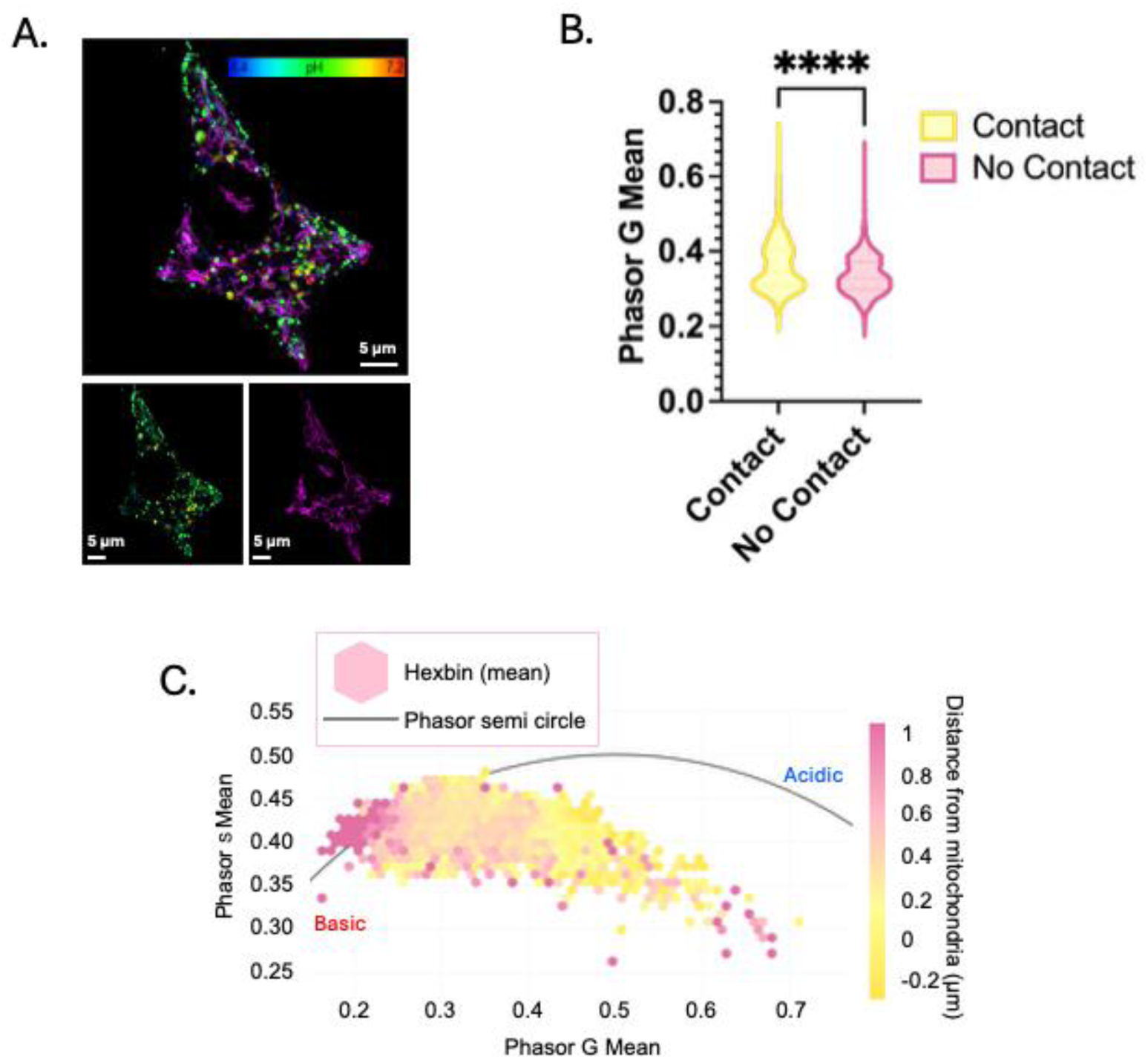
Mitochondrial proximity modulates ISG luminal acidity. (A) Representative confocal image of the mitochondrial network (magenta) and pH-labeled insulin secretory granules (ISGs), displayed as a pseudocolor pH map (4.4–7.2). Scale bars, 5 µm. (B) Violin plot showing the distribution of mean phasor G values for ISGs in contact with mitochondria (−0.355 to 0.05 µm from the mitochondrial surface) versus non-contact ISGs (0.05–1.0 µm away). Statistical comparison was performed using a Kolmogorov–Smirnov test (KS D = 0.1763, P < 0.0001). (C) Phasor plot of ISG fluorescence lifetime values colored by distance from mitochondria. Hexbins represent mean values per bin, and the semicircle indicates the universal phasor circle. Points positioned further left along the semicircle correspond to more acidic luminal pH, demonstrating a shift in acidity associated with mitochondrial proximity.

### ISG association with the mitochondria changes with ISG age

To determine how the spatial relationship between ISGs and the mitochondrial network evolves with granule age, we employed a pulse–chase labeling strategy using a previously developed INS cell line expressing a genetically encoded SNAP-tag on proinsulin (**Figure 4A**) (38). Prior studies suggest that ISGs remain in an immature state for up to 3 hours following biogenesis based primarily on radiolabeled measurements of proinsulin conversion (6, 39). However, such approaches do not necessarily capture ongoing changes in granule acidification, condensation, or metabolic remodeling that may extend past this 3-hour window. Consistent with this idea, our spatial analysis shows that 3-hour ISGs remain predominantly perinuclear and closely associated with the TGN (**Figure S3**), indicating that large-scale trafficking toward the cell periphery has not yet occurred. Radial redistribution increases between 6 and 12 hours (**Figure S4A**), suggesting that granule transport and maturation-associated remodeling extend beyond the initial proinsulin cleavage window. Given that trafficking and maturation are thought to be coordinated processes, these observations support the idea that functionally relevant maturation events persist past 3 hours.

**Figure 4.**
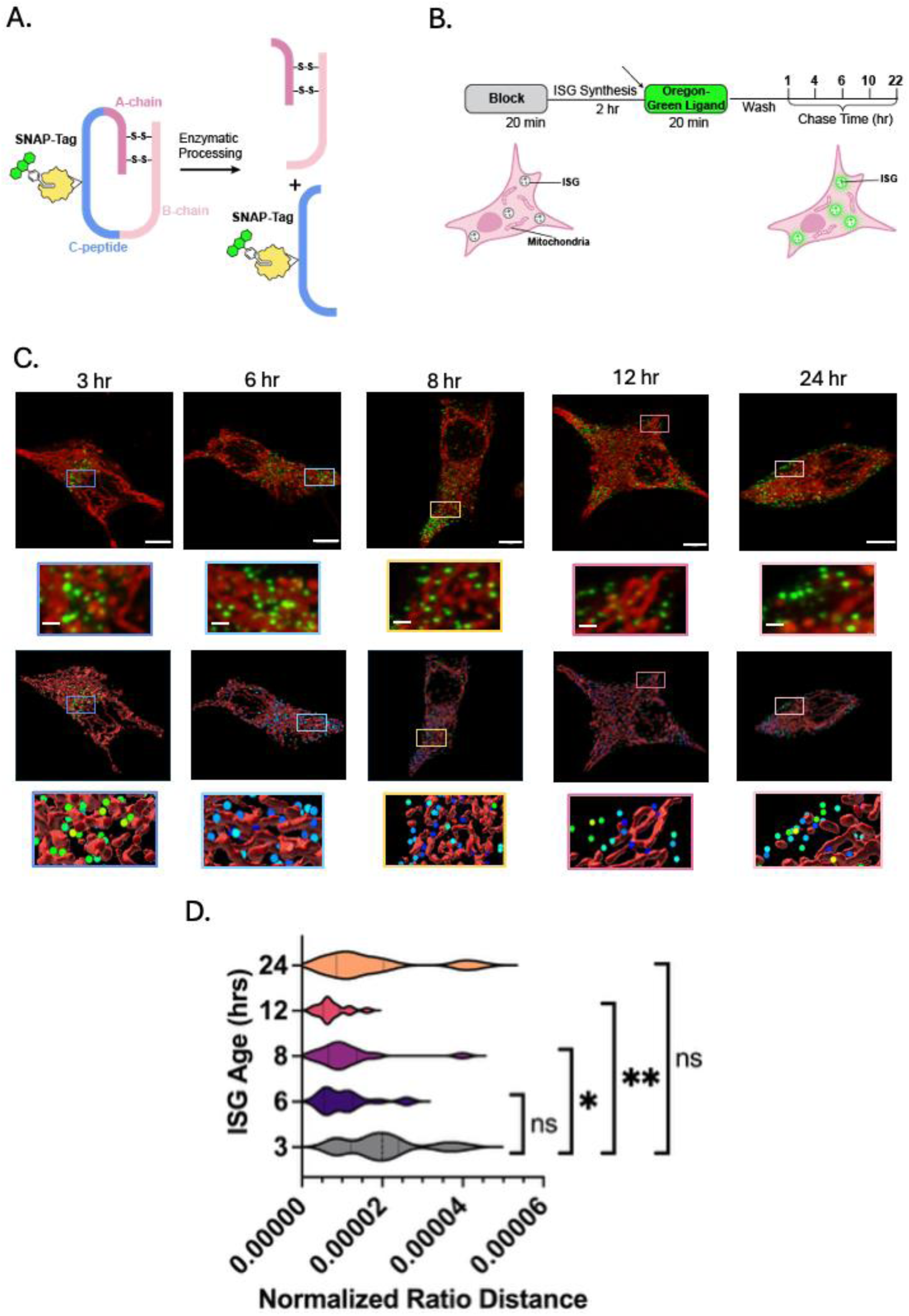
ISG age influences mitochondrial association. (A) Schematic of the pro–C-peptide–SNAP construct and its post-translational processing into insulin and C-peptide–SNAP. Diagram illustrating the fluorescent pulse–chase labeling strategy used to track ISG age. (B) Cells were pulse-labeled with SNAP–Oregon Green and chased for the indicated times prior to fixation. Two hours before fixation, cells were incubated with PK-Mito Orange to label the mitochondrial network. (C) Representative confocal images of whole cells (scale bar, 5 µm) and magnified insets (scale bar, 1 µm) are shown. Three-dimensional reconstructions were generated in Imaris to segment ISGs and mitochondria from fluorescence signals. (D) Normalized distance of ISGs to the nearest mitochondrion was quantified using Imaris-generated segmentation masks. For each cell, the fraction of ISGs located ≤0.1 µm from the nearest mitochondrion relative to total ISGs was calculated and normalized to the mitochondrial surface volume mask. Statistical significance was assessed using a Kolmogorov–Smirnov (KS) test with the 3-hour time point as the reference control: 3 h vs 6 h (KS D = 0.4286), 3 h vs 8 h (KS D = 0.5714, P = 0.0207), 3 h vs 12 h (KS D = 0.6429, P = 0.0061), and 3 h vs 24 h (KS D = 0.3571).

This later window remains relatively understudied but is likely critical for understanding how ISG maturation is regulated following synthesis. ISGs are known to interact with other organelles during their transit toward the cell periphery (6, 21, 22), we hypothesized that interactions with the mitochondrial network may contribute to maturation during these stages. We therefore focused our pulse–chase analysis on time points spanning and extending beyond the reported proinsulin conversion window, analyzing ISGs at 3, 6, 8, 12, and 24 hours of age (**Figure 4B**).

To quantify proximity to the mitochondria, we calculated for each cell, the fraction of ISGs located ≤0.1 µm from the mitochondrial network relative to the total ISG population and normalized this ratio to mitochondria volume to account for intercellular differences (**Figure 4D**). Younger (3 and 6 hour) old ISGs had more contacts with the mitochondria network, as described by a larger ratio. Notably, intermediate chase times (hours 8 and 12) exhibited a reduction in mitochondria-ISG proximity relative to the earlier tested time points, suggesting that mitochondria proximity is temporally regulated after initial proinsulin conversion. Analysis of absolute ISG–mitochondria distances without normalization yielded similar trends (**Figure S4B**), with increased separation at intermediate ages. Together, these findings indicate that mitochondrial proximity is dynamically modulated across the ISG lifetime and may persist beyond the initial 3-hour window defined by proinsulin cleavage, potentially supporting continued granule acidification and structural remodeling.

Our data reveal age-dependent differences in apparent ISG number (**Figure S4D**), with the 3-hour time point exhibiting significantly fewer detectable vesicles compared to later time points. This reduction may reflect limitations in confocal resolution, as newly formed ISGs are likely positioned in close proximity following biosynthesis, rendering individual granules difficult to resolve as distinct objects. To distinguish changes in granule number from differences in spatial organization, we next assessed ISG clustering directly.

Spatial clustering analysis has previously been used to characterize ISG spatial organization and identify distinct granule subpopulations (24, 40). To assess whether ISG localization varies with age, we quantified local ISG clustering by calculating the average distance to the three nearest neighboring ISGs for each granule, providing a measure of local vesicle packing relevant to trafficking and organization (**Figure S4E**). Distances were normalized to the population median at the 3-hour chase time to account for variability in absolute nearest-neighbor distances arising from differences in cell size and ISG number.

Intermediate chase times exhibited increased average nearest-neighbor distances relative to early chase conditions, indicating reduced local clustering, whereas early and late chase times displayed more compact spatial organization. Notably, the period of enhanced clustering (3–8 hours) temporally overlaps with the window during which ISGs exhibit shorter average distances to the mitochondrial network in our proximity analysis. While clustering and mitochondrial proximity represent independent measurements, the parallel age-dependent trends suggest that early ISG organization and mitochondrial association may be coordinated features of granule maturation.

## Discussion

Although ISGs have been previously observed in proximity to the mitochondrial network, the functional significance of these interactions has remained unclear. Our data now provide evidence that mitochondria-associated ISGs exhibit increased biomolecular density and enhanced acidification. The marked increase in density near mitochondria suggests changes in granule composition consistent with insulin condensation, while the observed reduction in luminal pH further supports progression along the maturation pathway.

Importantly, our temporal analyses indicate that this association is enriched after the initial ∼3-hour window commonly used to define proinsulin conversion. Because radiolabeled studies primarily report on peptide cleavage, they do not capture subsequent remodeling events. We therefore propose that mitochondrial association can occur after proinsulin processing and represents a post-cleavage phase of ISG remodeling characterized by continued acidification, structural reorganization, and compositional refinement.

In contrast, ISGs not associated with mitochondria generally lack these maturation-associated features, with the exception of granules located at or near the plasma membrane. These peripheral granules likely represent a secretion-competent population that has already completed earlier remodeling steps and therefore no longer requires mitochondrial engagement.

As suggested by our cryo-ET data, the close spatial proximity between ISGs and mitochondria, together with our observation of electron-dense material at their interface (**Figure S5**), indicates a potential functional coupling between these two organelles, likely mediated through proteins.

Lipid transfer from the mitochondria has been observed with other organelles (41–43), making it plausible that mitochondria transfer lipids to ISGs, potentially contributing to the increased LAC values observed in mitochondria-proximal granules. Alterations in lipid composition reshape membrane biophysical properties, including charge and curvature, which in turn affect the localization and activity of membrane proteins such as V-ATPases and other transporters. These changes can regulate ion flux across the granule membrane and influence the acidification and condensation processes that define ISG maturation.

Notably, immature and mature ISGs exhibit distinct lipid compositions, with older granules enriched in phosphatidylethanolamine (PE) relative to younger vesicles (28). Mitochondria are rich in specific lipids, including PE (44, 45). This overlap raises the possibility that mitochondrial association contributes to the accumulation of PE in maturing ISGs. Supporting this idea, VPS13C, a lipid transfer protein implicated at organelle contact sites involving the mitochondria (41), has been shown to transport PE (41, 46).

Consistent with this possibility, our cryo-ET resembles the structural features reported for VPS13 family lipid transfer proteins (41). While we cannot assign molecular identity based on morphology alone, this similarity supports the hypothesis that mitochondria–ISG contacts may facilitate PE transfer via a VPS13C-like mechanism, thereby promoting membrane remodeling during ISG maturation. Future work employing advanced cryo-ET denoising and molecular identification approaches will be required to define the identity and directionality of lipid transfer at these sites (47).

In addition to being rich sources of lipids, mitochondria create a high local concentration of ions such as zinc and calcium, as well as ATP and glutamate (48–55). Ca^2+^ and Zn^2+^ ions are essential for proper ISG function and cargo processing (7, 8). Elevated local concentrations likely enhance energy-dependent transport and ion flux into ISGs, thereby promoting their maturation and functional competence. Another critical step in insulin maturation is acidification of the granule lumen, which is mediated by V-ATPase proton pumps located on the ISG membrane (33, 56, 57). Because these proton pumps are ATP-dependent, proximity to the mitochondrial network, where ATP concentrations are locally elevated may facilitate efficient granule acidification. Consistent with this idea, our data show that ISGs positioned near mitochondria exhibit increased acidity.

In addition, the mitochondrial network increases the local concentration of glutamate (54, 55). ISGs express glutamate transporters on their membranes (58), and glutamate functions as a counterion for H+, enabling sustained V-ATPase activity and continued luminal acidification (59, 60). Thus, the localized availability of ATP and glutamate near the mitochondrial network may promote the acidification necessary for proinsulin processing and the progression of insulin maturation within immature granules.

SXT analysis reveals that the majority of mitochondria–ISG contacts are localized within the cell interior, consistent with our pulse–chase observations showing that 3-6-hour-old granules exhibit mitochondrial proximity in these regions. Notably, ISGs located in the cell interior and in contact with mitochondria exhibit increased acidity, aligning with prior reports that centrally positioned granules tend to be more acidic (37). Although we also observe acidic granules in the interior that do not appear to contact mitochondria, their morphology suggests that these may represent recycling ISGs rather than actively maturing granules.

Our pulse–chase analysis indicates that mitochondria–ISG contacts are enriched during the younger (3-6 hour) window after synthesis. These contacts decrease at intermediate ages (8-12 hours) and increase again in late stage (24 hour) ISGs. Previous studies have determined insulin maturation to take on average 3 hours (6, 39). These studies are based primarily on proinsulin conversion to insulin. The timing of contact enrichment observed here therefore falls within a maturation window after initial proinsulin to insulin conversion, raising the possibility that mitochondrial interactions contribute to late-stage granule remodeling rather than being restricted to the earliest phases of ISG biogenesis.

Together, our data support a working model in which ISG trafficking and maturation are spatially coordinated around the mitochondrial network. Following synthesis, ISGs are positioned in a perinuclear region near the trans-Golgi network. As granules age, our pulse–chase analysis indicates that they redistribute radially toward the cell periphery while establishing contacts with mitochondria, potentially to support maturation-related processes.

Collectively, these findings suggest that mitochondrial interactions may facilitate maturation within a distinct subpopulation of ISGs and accommodates the presence of mature granules that lack detectable mitochondrial contact. Such mitochondria-associated ISGs may therefore represent a functionally specialized pool that undergoes more rapid maturation, potentially priming them for timely insulin release in response to metabolic demand.

In this context, mitochondrial dysfunction, commonly observed in diabetes, may have important consequences for ISG maturation. Several studies have reported impaired mitochondrial function in diabetic β-cells, including increased reactive oxygen species production, reduced ATP generation and increased mitochondrial fragmentation (61–63). If mitochondrial proximity supports efficient granule acidification and insulin processing, then compromised mitochondrial metabolism could attenuate this facilitative effect, potentially leading to dysfunctional insulin maturation.

Despite the insights provided here, several important questions remain regarding the molecular composition and precise function of mitochondria–ISG contact sites. Defining the mechanisms that establish and regulate these interactions will be essential for understanding how they may become dysregulated in metabolic disease. In addition, refining the temporal resolution of pulse–chase experiments with shorter labeling intervals may better define when these interactions arise during the maturation timeline. Coupling age-defined subpopulations with biochemical readouts, such as intragranular pH, could further resolve the functional state of granules engaged with mitochondria and illuminate the metabolic processes occurring at these sites.

There are several technical considerations that have an impact on our conclusions. Fluorescence microscopy is limited by diffraction, which constrains the precision with which mitochondria–ISG contacts can be resolved and quantified in segmentation analyses. Moreover, chemical fixation may introduce artifacts that perturb native cellular organization. Although the integration of multiple imaging modalities allowed us to examine these interactions across scales, differences in sample preparation and imaging conditions can alter cellular morphology, potentially influencing subcellular spatial analyses.

In summary, our findings support a model in which insulin granule maturation is not solely a temporally defined process, but also a spatially organized one. By integrating multi-scale imaging approaches, we identify mitochondria–ISG proximity as a potential facilitator of granule acidification, condensation, and spatial remodeling. These observations expand our understanding of organelle neighborhoods in β-cells and provide a framework for investigating how metabolic dysfunction may impair insulin production in diabetes. Future efforts to define the biochemical mechanism of this contact will be critical for determining whether they represent regulated contact sites analogous to other well-characterized organelle contacts, as well as how this contact impacts granule maturation.

## Acknowledgements

We graciously thank Samuel Stephens at the University of Iowa for gifting the SNAP-Tag INS cell line. We thank Valentina Loconte and Kevin Chang for their contributions to SXT data acquisition and analysis, respectively. Thanks to Ashley Archambeau and Htet Khant for Cryo-ET sample prep and data acquisition. Thanks to the Core Center of Excellence in Nano Imaging (CNI), the Translational Imaging Core (TIC), and the National Center for X-Ray Tomography (LBNL, Berkeley) for use of the Thermo Glacios and Titan Krios, Leica SP8 Lightning confocal microscope, and the XM-2 soft X-ray microscope, respectively. Thanks to Janille Cuala for supplying the genetic mouse lines. We are grateful to members of the Kate White and Vadim Cherezov labs, and to the reviewers for their feedback on this work. We are thankful to the Cell Culture Core (USC) and the Bridge Structural Biology Center (SBC), particularly Jeffrey Velasquez, for help in maintaining cell stocks and molecular biology experiments, respectively. We would also like to thank Katya Cherezov for designing the SNAP-Tag workflow diagrams in figure 2. Funding from the National Institute of General Medical Sciences of the National Institutes of Health under award number R35GM154893 and the Bridge Institute at USC helped support this work.

## Author Contributions

R.K. and K.L.W. conceived the project with contributions from all authors on experimental design, data interpretation, and manuscript editing. R.K and W.L. collected pH data, A.D. collected SXT data. All pulse-chase data collected was done by R.K. Blender renders were done by R.V. All authors have given approval to the final version of the manuscript.

## Materials and Methods

### Cell culture and reagents

ProCpepSNAP expressing INS-832/3 cells (A gift from Dr. Samuel Stephens, University of Iowa, IA, USA) were cultured in optimized RPMI 1640 (AddexBio, cat#C0004-02) containing 10% FBS (Cytiva, Cat#83007) and 0.05mM β-Mercaptoethanol (Thermo Scientific, Cat#21985023) at 37°C and 5% CO_2_. Cells were seeded at a density of 4.0 x 10^4^ cells/cm^2^ in T75 flasks and passaged in accordance with previously documented protocols (40).

Rat insulinoma INS-1E cells (obtained from Pierre Maechler, University of Geneva) were cultured in optimized RPMI supplemented with 5% FBS at 37°C and 5% CO_2_. INS-1E cells were plated at a density of 4.0 x 10^4^ cells/cm^2^ in T75 flasks and passaged in accordance with previously documented protocols (41).

For imaging and pulse-chase experiments, either INS-1E or ProCpepSNAP cells were plated or polymer chambered coverslips (Ibidi, cat#80806). For transfections, INS-1E cells were initially seeded at a density of 40,000/cm^2^ and grown for 48 hours to ensure adherence and confluency prior to transfection. Lipofectamine 3000 reagents (Thermo Fisher Scientific, cat# L3000001) were used as per the instructions provided by the manufacturer.

### Animal Methods

For cryo-ET experiments involving primary β-cells, animal procedures were approved and conducted per the Institutional Animal Care and Use Committee (IACUC) guidelines at the Children’s Hospital of Los Angeles (Animal Use Protocol #337). The LSL-Salsa6f mouse model expressing expressing tdTomato linked to GCaMP6f by a V5 epitope tag was created by crossing B6(Cg)-Ins1tm1.1(cre)Thor/J (JAX #026801) with B6(129S4)-Gt(ROSA)26Sortm1.1(CAG-tdTomato/GCaMP6f)Mdcah/J (JAX# 031968).

### SXT sample preparation

INS-1E cells were starved for 30 minutes at 37°C in a Krebs-Ringer bicarbonate-based buffer (KRBH; 115 mM NaCl, 24 mM NaHCO_3_, 5 mM KCl, 1 mM MgCl_2_, and 1 mM CaCl_2_), additionally supplemented with 0.2% bovine serum albumin (BSA), 1.1mM glucose, and 10nM HEPES maintained at a pH of 7.4. Post stimulation, cells were washed with 1XPBS, collected, and resuspended in 1XPBS and kept on ice. Following this, cells were inserted into microcapillaries and plunge frozen into liquid ethane to cryo-preserve the native state of the cell.

### SXT data collection and Tomogram segmentation

SXT was conducted at the National Center for X-Ray Tomography (Lawrence Berkeley National Laboratory, Berkeley, CA) using the XM-2 microscope. Imaging was performed at the “water window” energy range. Post imaging, projection images were reconstructed into 3D tomograms and segmented using Amira (Thermo Fisher Scientific). Organelles were identified and segmented based on previous reports (21, 26, 27). Detailed information about imaging and segmentation parameters can be found in Deshmukh et al. (23)

### SXT Organelle quantification and data analysis

Post-segmentation, ISG and mitochondrial organelle masks were used for further analysis. The distance between ISGs and mitochondria was calculated using custom Python code measuring the shortest distance between the mitochondrial surface and any ISG voxel. ISGs were then classified into “contact” or “no contact” classes based on a 50 nm cut-off. ISG localization was calculated by generating a Euclidean distance transform (EDT) map for each cell. Normalization was performed by dividing the mean EDT value of each ISG by the maximum EDT value of the cell in which it resided. This normalized EDT value was then used to compare localization between classes. ISG diameter and LAC were calculated similarly to previous reports, as described in detail in Deshmukh et al. (23)

### Confocal Microscopy

Unless otherwise noted, all imaging experiments were performed on a Leica SP8 DIVE FALCON with a 63x/1.2NA water immersion objective. For pulse-chase ProCpepSNAP experiments, the following excitation and emissions were used: Oregon-Green SNAP 488 (Ex 490nm; Em 514-550nm), PK-Mito Orange (Ex 591nm; Em 600-700nm).

### FLIM pH imaging

Live INS1E cells expressing NPY-E2GFP were stained with pKmito Deep Red according to the manufacturer’s protocol. Three-dimensional z-stack images were acquired on a Leica Stellaris 8 tau-STED microscope using an 86×/1.2 NA water-immersion objective (512 × 512 pixels; field of view containing a single cell). Two channels were acquired sequentially in photon-counting mode: green (Ex 473 nm; Em 497–519 nm) and red (Ex 653 nm; Em 666–829 nm). FLIM was enabled in the green channel for pH measurements using 80 MHz pulsed excitation. Photons were accumulated over 4 scans, and the final image was binned 2 × 2 for FLIM analysis. FLIM data were analyzed using phasor analysis, and pH values were obtained using the calibration/relationship reported previously (46).

### Cryo-ET Sample Preparation

Cryo-ET imaging was performed on primary mouse beta cells. Briefly, the LSL-Salsa6f mouse model expressing tdTomato linked to GCaMP6f by a V5 epitope tag, to indicate beta-cell identity and calcium signaling, respectively, was used for islet isolation. Islets were dissociated into single cells using previously established protocols (42). Lacey carbon grids were used for all experiments. Grids were first sterilized with 70% ethanol and dried in 35mm plates equipped with coverslips on the bottom. After drying, grids were incubated with poly-D-lysine. Post-incubation, grids were washed three times with 1XPBS, after which the dissociated single cells were added. Additional media was added to sustain the cells, and the grids were then incubated overnight at 37°C and 5% CO_2_. After overnight incubation, grids were screened for cell quality using brightfield imaging. The best grids were then selected for further processing. Next, grids were plunge frozen in liquid ethane, clipped into autogrids, and stored in liquid nitrogen until imaging.

### Cryo-ET Imaging and Processing

Autogrids were screened under cryogenic conditions using a 200kV Glacios cryo-TEM equipped with a Falcon4 detector (Thermo Fisher Scientific; magnification 13,500X or 93,000X). Selected samples were then chosen for further cryo-ET imaging. Tilt series were collected on a 300kV Krios G3i equipped with a Gatan K3 direct detection camera (Thermo Fisher Scientific; 26,000X magnification). Tilt series were acquired in increments of 3 degrees from -60° to +60° in a dose-symmetric fashion. Individual tilt series were recorded with a defocus range of -4 to -6 µm. The cumulative dose for each tilt series was between 80 and 120e/Å^2^. The collected tilt series were reconstructed into 4X binned tomograms using IMOD, through SIRT-based reconstruction with seven iterations.

### Pulse-chase SNAP-tag labeling and Imaris segmentation

ProCpepSNAP expressing 832/3 cells were plated onto polymer chambered coverslips at a density of 30,000 cells/cm^2^ in normal growth media and grown overnight at 37°C and 5% CO_2_. For SNAP-labeling, cells were initially labeled with SNAP cell block (10µM, NEB) diluted in normal growth media for 30 minutes at 37°C and 5% CO_2_. The cells were washed 3 times for 10 minutes each in normal growth media and cultured in normal growth media for 2 hours. Cells were then labeled with SNAP Oregon-green (1 µM, NEB) diluted in normal growth media at 37°C and 5% CO_2_ before being washed 3 x 10 minutes in reduced glucose (5mM) growth media. were chased in reduced glucose media for the indicated times prior to fixation. 2 hours prior to fixation, cells were labeled with PKmito Orange FX (500nM, SpiroChrome) in reduced glucose media.

Fixation was performed by doing a 20 second prefixation in pre-warmed 2% glutaraldehyde at room temperature, followed by fixation with 4% paraformaldehyde for 8 minutes at room temperature. Cells were washed 3 times for 5 minutes each with 1X PBS and stored in 1X PBS at 4°C protected from light prior to imaging.

Mitochondria and ISG segmentation were completed using Imaris 10.2.1 software. Segmentation for ISGs was done using the following parameters: Enable Region Growing = false, Enable Tracking = false, Enable Classify = false, Enable Region Growing = false, Enable Shortest Distance = true, XY Diameter = 0.350 µm, Estimated Z Diameter = 0.700 µm, Background Subtraction = true. Intensity threshold was manually adjusted to capture

ISGs based on judgment per cell. Segmentation for mitochondria was done using the following parameters: Region Growing = false, Enable Tracking = false, Enable Classify = false, Enable Shortest Distance = true, Enable Smooth = true, Surface Grain Size = 0.100 µm, Enable Eliminate Background = true, Diameter Of Largest Sphere = 0.120 µm. Intensity threshold was manually adjusted per cell to include 80% of signal.

### Statistical Analysis

All quantitative data were graphed and analyzed using GraphPad Prism 10.3.0 software. Statistical analysis was performed within Prism and is described in the figure legends along with sample size for each analysis. Statistical significance was set at *p < 0.05; **p < 0.01; ***p < 0.001; ****p < 0.0001 for all analyses.

