## Supplemental_Figures for "Integrated biophysical and spatial remodeling during insulin secretory granule maturation at the mitochondrial network"

For

### 6 Supplementary Figures

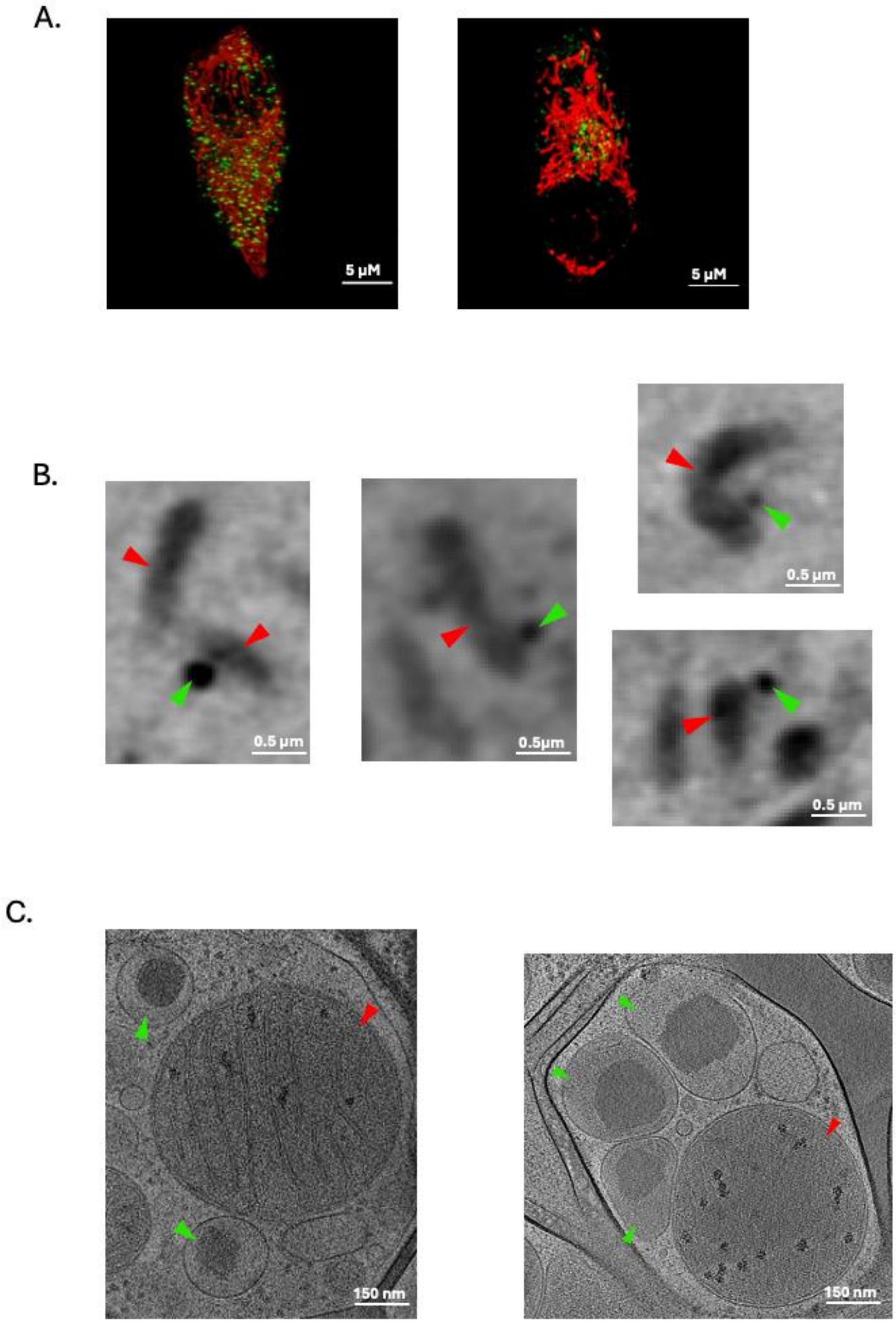


**Supplementary Figure 1. Additional examples of ISG–mitochondria contacts visualized by correlative fluorescence, soft X-ray tomography, and cryo-electron tomography.**

(A) Representative confocal fluorescence images showing insulin secretory granules (ISGs, green) in proximity to the mitochondrial network (red). Scale bars, 5 μm. (B) Soft X-ray tomographic slices highlighting close apposition between mitochondria (red arrowheads) and ISGs (green arrowheads). Scale bars, 0.5 μm. (C) Cryo-electron tomographic slices confirming membrane-bound ISGs (green arrowheads) closely apposed to the outer mitochondrial membrane (red arrowheads) without evidence of membrane fusion. Scale bars, 150 nm.

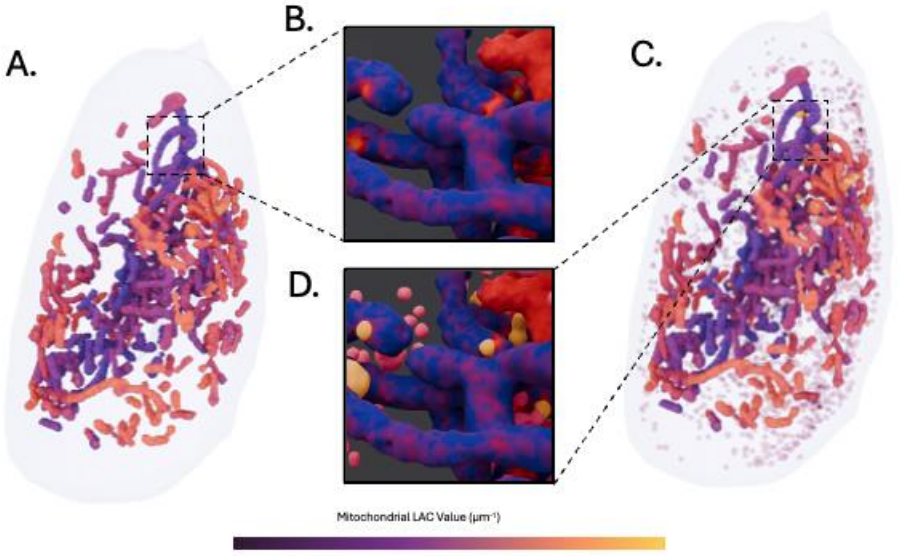


**Supplementary Figure 2. Local variation in the biomolecular density (LAC) of mitochondria next to ISGs-mitochondria contact sites**

(A) Whole cell rendering of a 3D mitochondrial network in a representative INS-1E cell showing each mitochondrial voxel colored based on its LAC value (Purple – Low LAC; Yellow – High LAC). (B) Close-up view of the inset in A showing localized LAC hotspots on mitochondria. (C) Whole cell rendering of a 3D mitochondrial network (similar to (A)) with ISGs not in contact with mitochondria (Transparent) and ISGs in contact with mitochondria (Yellow). (D) Zoomed in view of the inset in C showing that the ISG in contact with mitochondria are localized to the LAC hotspots shown in (B).

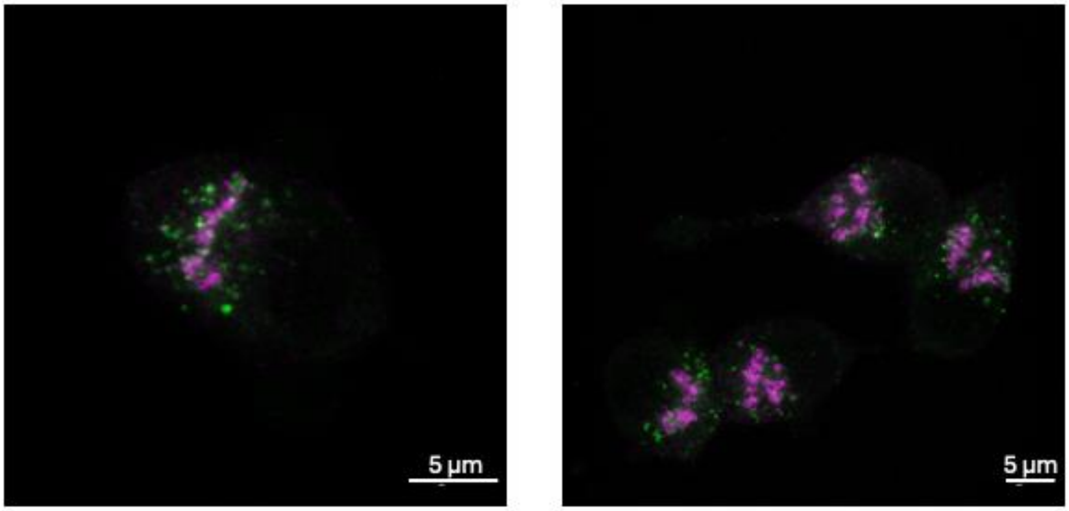


**Supplementary Figure 3. Representative fluorescence images of ISGs and TGN**

Confocal images showing insulin secretory granules (ISGs, green) and the TGN (magenta) in independent cells. Scale bars, 5 µm.

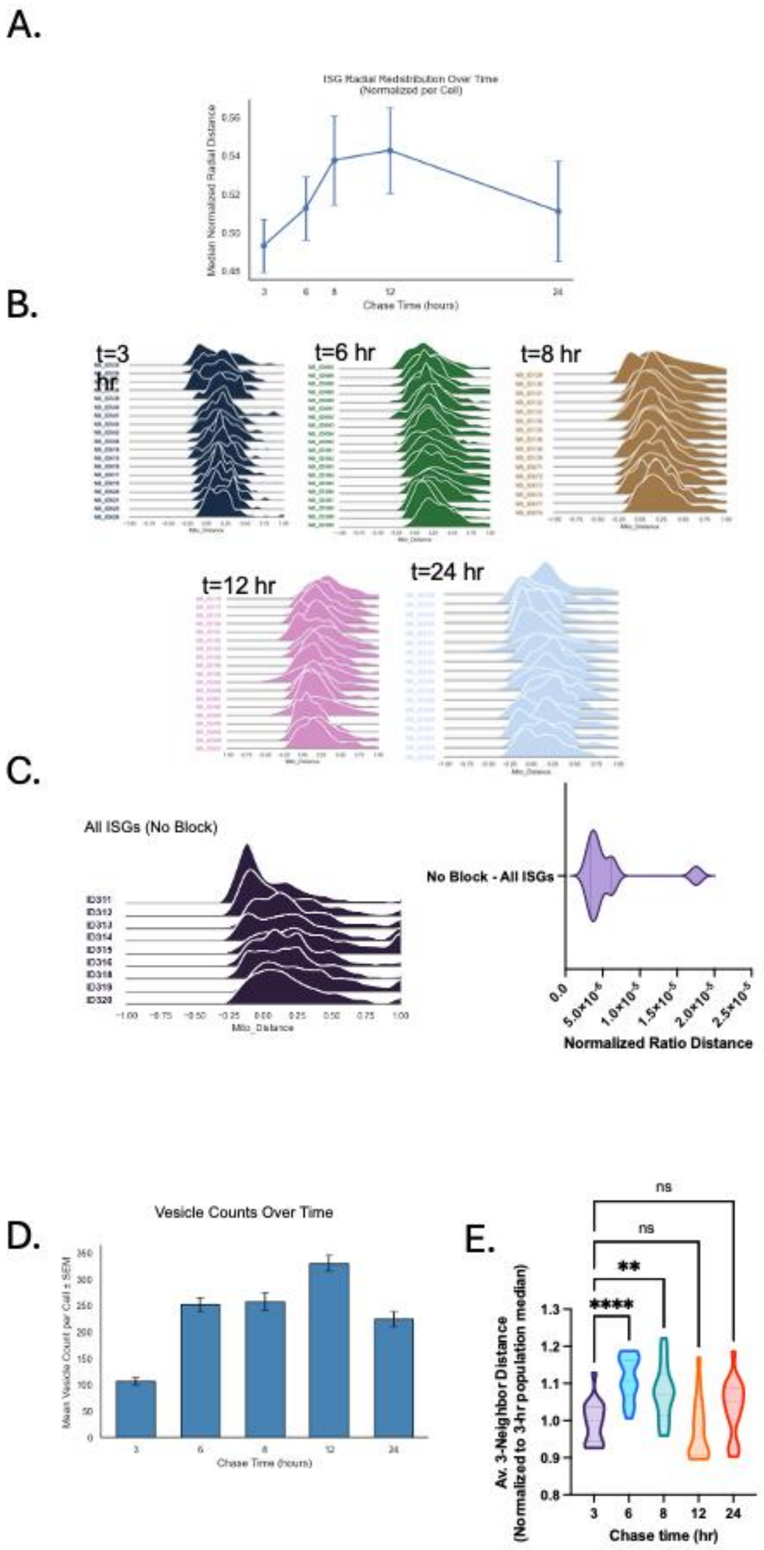


**Supplementary Figure 4. Age-dependent radial redistribution and abundance of ISGs during pulse–chase labeling.**

(A) Median normalized radial distance of ISGs from the cell center over chase time (3–24 h), normalized per cell. Data are shown as mean ± SEM. (B) Ridge plots showing the distribution of ISG distances to the nearest mitochondrion for individual cells at each chase time point (1, 4, 6, 10, and 22 h). Each curve represents a single cell. Negative values indicate closer proximity to mitochondria, whereas positive values indicate increasing distance. (C) Representative distribution of ISG–mitochondria distances in non-block conditions, displayed per cell. (D) Bar plot shows the mean number of ISGs per cell at each chase time, with error bars indicating SEM. While ISG count changes over time, these differences do not necessarily align with the observed shift in ISG-mitochondria neighbor distances, indicating that clustering behavior may change as a function of ISG age.

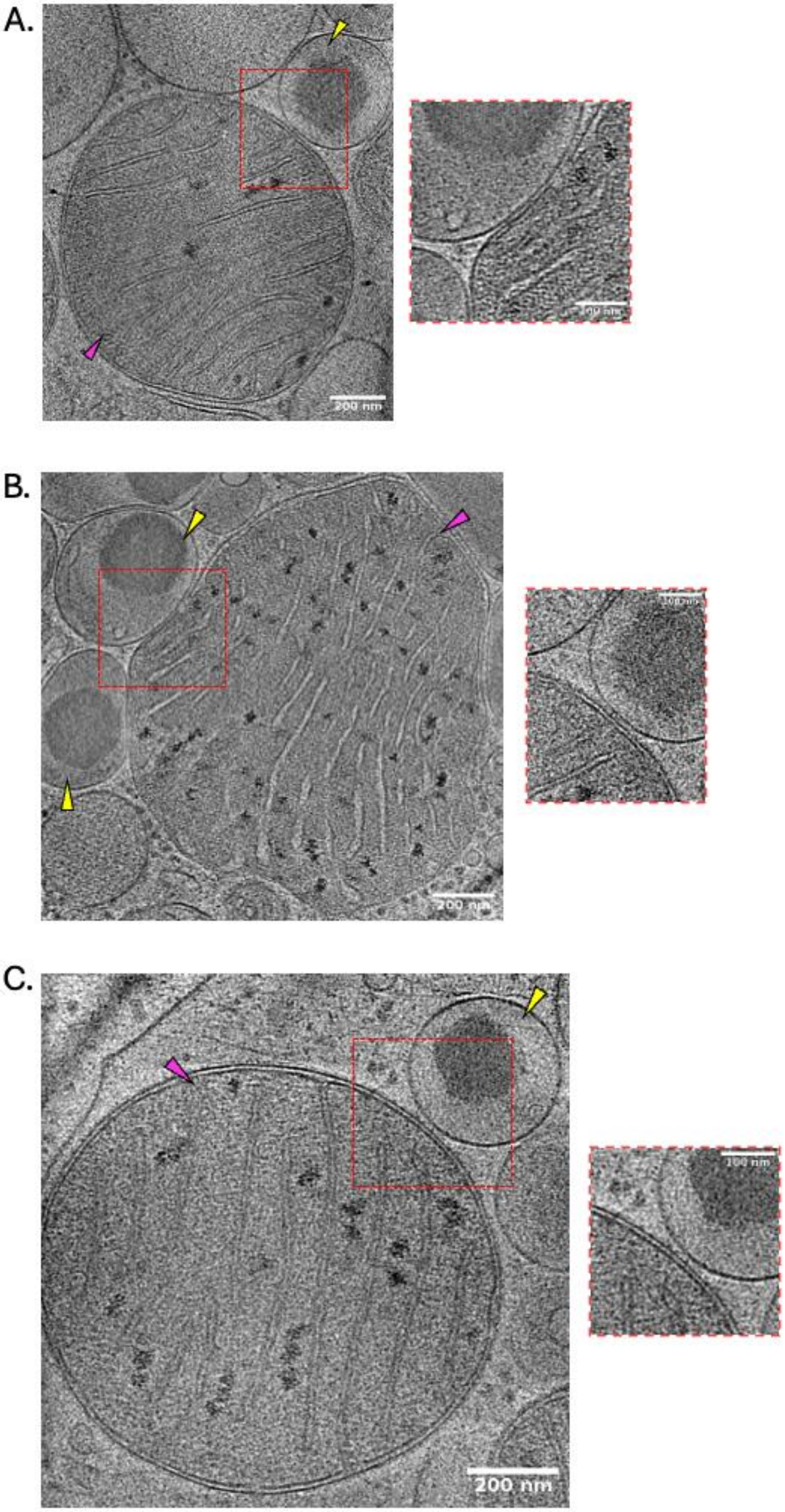


**Supplementary Figure 5. Cryo-electron tomography reveals tight ISG–mitochondria contacts with putative inter-organelle densities**

(A–C) Representative cryo-ET slices showing insulin secretory granules (ISGs; yellow arrowheads) in extremely close apposition to mitochondria (pink arrowheads). Red dashed boxes indicate regions enlarged at right. Insets highlight narrow membrane separation distances and electron-dense material at the interface, consistent with potential protein structures between the ISG membrane and the outer mitochondrial membrane. Scale bars, 200 nm (main panels) and 100 nm (insets).
